# Different temporal trends in vascular plant and bryophyte communities along elevational gradients over four decades of warming

**DOI:** 10.1101/2020.03.17.994814

**Authors:** Antoine Becker-Scarpitta, Diane Auberson-Lavoie, Raphael Aussenac, Mark Vellend

## Abstract

Despite many studies showing biodiversity responses to warming, the generality of such responses across taxonomic groups remains unclear. Very few studies have tested for evidence of bryophyte community responses to warming, even though bryophytes are major contributors to diversity and functioning in many ecosystems. Here we report an empirical study comparing long-term change of bryophyte and vascular plant communities in two sites with contrasting long-term warming trends, using “legacy” botanical records as a baseline for comparison with contemporary resurveys. We hypothesized that ecological changes would be greater in sites with a stronger warming trend, and that vascular plant communities, with narrower climatic niches, would be more sensitive than bryophyte communities to climate warming. For each taxonomic group in each site, we quantified the magnitude of changes in species’ distributions along the elevation gradient, species richness, and community composition. We found contrasted temporal changes in bryophyte vs. vascular plant communities, which only partially supported the warming hypothesis. In the area with a stronger warming trend, we found a significant increase of local diversity and beta-diversity for vascular plants, but not for bryophytes. Presence absence data did not provide sufficient power to detect elevational shifts in species distributions. The patterns observed for bryophytes are in accordance with recent literature showing that local diversity can remain unchanged despite strong changes in composition. Regardless of whether one taxon is systematically more or less sensitive to environmental change than another, our results suggest that vascular plants cannot be used as a surrogate for bryophytes in terms of predicting the nature and magnitude of responses to warming. Thus, to assess overall biodiversity responses to global change, abundance data from different taxonomic groups and different community properties need to be synthesized.

## Introduction

Ecological impacts of global changes on biodiversity can take many different forms (McGill et al. 2015, Vellend et al. 2017b). In the context of climate warming, montane ecosystems have received considerable attention due to clear predictions of changes in species’ distributions (upslope shifts) and community composition (favoring warm-adapted species) along elevation gradients. The many studies to date have focused on relatively few taxonomic groups, in particular vascular plants, invertebrate and vertebrates (Lenoir et al. 2008, Chen et al. 2011, Pauli et al. 2012, Stockli et al. 2012, Rumpf et al. 2018, 2019), and there is now a need for more exhaustive multi-taxonomic syntheses (Lenoir and Svenning 2015, Lenoir et al. 2020). The few studies of bryophyte responses to climate warming suggest that results might be different than those for vascular plants (Walker et al. 2006, Bergamini et al. 2009, Hudson and Henry 2010, Raabe et al. 2010, Vanneste et al. 2017, Becker-Scarpitta et al. 2017).

Understanding variation among taxonomic groups in their responses to environmental change is crucial for identifying priorities in conservation. However, the spatial variability of climate change complicates comparisons between studies of different taxa in different regions, and it is difficult to predict *a priori* differences between taxonomic groups. Thus, our limited knowledge base with which to identify the most relevant set of metrics or the taxonomic groups most sensitive to environmental change constrains our ability to set efficient conservation priorities. We operationally define “sensitivity” here as responsiveness: i.e., the degree to which a given community property changes in the face of environmental change.

To assess long-term responses of ecological communities to environmental change, “legacy” ecological records have been used to provide baselines for comparison with contemporary resurveys (Vellend et al. 2013, Chytrý et al. 2014, Verheyen et al. 2017, Hédl et al. 2017). However, historical botanical studies are strongly biased towards vascular plants, with few data on bryophytes, due in part to the difficulty of identification (Gignac 2001; He, He, and Hyvönen 2016; Möls et al. 2013 but see Becker-Scarpitta et al., 2017; Bergamini et al., 2009; Delgado & Ederra, 2013; Outhwaite, Gregory, Chandler, Collen, & Isaac, n.d.; Vanneste et al., 2017). This is despite the fact that bryophytes are significant contributors to diversity in many temperate ecosystem, and a major component of the vegetation cover in boreal or alpine ecosystems. Bryophytes play a crucial role in ecosystem functions, such as biomass accumulation, water retention, nutrient cycling, and food web dynamics (Turetsky 2003, Rydin 2008, Lindo and Gonzalez 2010, Tuba et al. 2011, Patiño and Vanderpoorten 2018).

For several reasons we might expect vascular plants and bryophytes to show different responses to various sources of environmental change (Lee and La Roi 1979, Möls et al. 2013, Bagella 2014, Vanneste et al. 2017, Becker-Scarpitta et al. 2017). Compared to vascular plants, bryophytes are distinguished by their small size, high sensitivity to the moisture and chemistry of their immediate microenvironment (i.e., they are poikilohydric), lower temperature optima for growth, absence of roots and an efficient vascular system, and dispersal via spores (Glime 2007). These two groups can show contrasted spatial patterns of diversity (Lalanne et al. 2008, Mateo et al. 2016). For instance, vascular plants show a clear latitudinal diversity gradient of decreasing species richness with increasing latitude, while this is not true for bryophytes, for which temperate latitudes are equally diverse as tropical latitudes (Geffert et al. 2013, Mateo et al. 2016). Some studies have also observed different patterns of community β-diversity (Kraft et al. 2011, Mateo et al. 2016) with vascular plant communities sometimes showing higher β-diversity along elevation gradients than bryophytes, suggesting a broader tolerance of bryophyte species to temperature (Lee and La Roi 1979, Vittoz et al. 2010, Glime 2013, Vanneste et al. 2017) or differences in dispersal ability (Lenoir et al. 2012). General predictions for the effects of warming on vascular plants include declines in the abundance of cold-adapted species, an upward expansion of elevational range limits for warm-adapted species (Rumpf et al. 2018), and an increase of local species richness in area without water-stress (Vellend et al. 2017b, Steinbauer et al. 2018, Harrison 2020). Compared to vascular plants, some studies have suggested that changes in bryophyte communities are more strongly influenced by stochastic processes or by micro-environmental variation than macro-environmental conditions (Pharo and Vitt 2000, Raabe et al. 2010, Fenton and Bergeron 2013). Because bryophytes have wider temperature tolerances and higher affinities to micro-environment than macro-environment, we might expect bryophytes to show lower sensitivity to global warming than vascular plants. The consequences of warming for β-diversity are more difficult to predict given a paucity of studies on this topic (Socolar et al. 2016 but see Nascimbene and Spitale 2017). However, numerical simulations show that species with high dispersal capacity will be favored in areas experiencing strong environmental changes, in which case we might predict a decrease of β-diversity (Mouquet and Loreau 2003) and thus biotic homogenization, at least temporarily (Clavel et al. 2011).

As in many parts of the world, eastern Canada has shown a general warming trend over the past ∼50 years, but with a strong east-west gradient in the magnitude of warming in the province of Québec (Yagouti et al. 2008). Becker-Scarpitta, Vissault, and Vellend (2019) showed that the magnitude of temporal changes of vascular plant communities in three protected areas generally increased from east to west in southern Québec, with greater changes in areas of stronger warming in recent decades. For two of these three parks, the historical data also included bryophytes, thus presenting an opportunity to test for differential sensitivity among taxonomic groups to warming. Forillon National Park is located at the eastern tip of the province of Québec where warming has been less pronounced than in Mont-Mégantic Provincial Park, which is in central Québec where the warming trend has been stronger (Savage & Vellend, 2015; Yagouti et al., 2008 & Supporting Information 1).

Here we report one of the first studies comparing long-term change of bryophytes and vascular plants communities in sites with contrasting long-term warming trends. In each of the two parks, we revisited ∼50 legacy vegetation plots initially surveyed in the 1970s, applying the same methods as the original surveys. To minimize potentially confounding factors, plots were selected in mature forest ecosystems that have not experienced major natural or anthropogenic disturbances during the time between surveys. We test two main hypotheses: (**i**) For both bryophytes and vascular plants, the park with a stronger warming trend (Mont-Mégantic) has experienced greater long-term community changes than the park with a weaker warming trend (Forillon); (**ii**) Vascular plant communities are more sensitive than bryophyte communities to climate warming. For each taxonomic group in each park, we quantified the magnitude of changes in (a) species’ distributions along the elevation gradient, (b) species richness (alpha-diversity), and (c) temporal change in community composition. Note that these predictions were already tested for vascular plants in Becker-Scarpitta et al. (2019), to which we add data for bryophytes in this paper.

## Methods

### Study site

Our two study sites were located in eastern Canada: Forillon National Park in eastern Québec (48°54′N, 64°21′W), and Mont-Mégantic Provincial Park in central Québec (45°27′N, 71°9′W). Neither park has experienced logging or forest management over the last ∼40 years. Forillon National Park was created in 1970 and covers 245 km^2^; our study plots ranged in elevation from ∼50 to 500 m a.s.l. The vegetation at Forillon is dominated by a mixture of northern tree species such as *Abies balsamea* (L.) Mill., *Picea glauca* (Moench) Voss and *Betula papyrifera* Marsh. at higher elevations, and *Acer saccharum* Marsh. and *Betula alleghaniensis* Britt. at lower elevations (Majcen 1981). Mont-Mégantic Provincial Park was created in 1994 (logging ceased in the 1960s during park planning) and covers ∼55 km^2^. Our study plots span an elevational gradient from ∼460 and 1100 m a.s.l., along which the vegetation gradient strongly resembles that in Parc Forillon: from temperate deciduous forests dominated by *Acer saccharum*., *Fagus grandifolia* Ehrh. and *Betula alleghaniensis*, to boreal forest dominated by *Abies balsamea*, and *Picea rubens* Sargent, Silva. (Marcotte and Grandtner 1974). Average temperatures at Mont-Mégantic shows an increase of ∼0.90°C between 1960 and 2005 while at Forillon the estimated increase is about ∼0.55°C for the same period (Appendix S1).

### Data set

Original vegetation surveys at both sites were conducted using phytosociological methods (Marcotte and Grandtner 1974, Majcen 1981). In each plot, authors listed all species in different strata (canopy trees, shrubs, herbs and ground bryophytes) and each species was assigned an abundance coefficient following Braun-Blanquet et al. (1952). For vascular plants, our analyses focused on shrubs and herbs, which were combined into a single “understorey” stratum. All bryophyte species that were found on the ground were recorded (i.e., on organic litter and soil surface mineral layers, not including obvious deadwood, tree trunks and rocks); these surveys did not involve intensive searches for individual stems of rare species within moss carpets (i.e., some locally rare species were missed). After consulting with botanists active in Québec in the 1960s and 1970s (C. Ansseau, Z. Majcen, personal communication, 2014), there remained uncertainty in the definition of the area over which percent cover was evaluated for bryophytes (microhabitats vs. entire plots). Thus, we are not confident in comparing Braun-Blanquet indices across time for bryophytes, as was done for vascular plants in Becker-Scarpitta et al. (2019). To maximize comparability across time and taxonomic group, we used presence-absence data for both vascular and bryophyte species in all statistical analyses.

Our approach to plot relocation is described in Becker-Scarpitta et al., (2019). In short, original survey plots were not permanently marked, although locations were reported in maps and/or tables, such that plots are considered “semi-permanent”. In both parks, original surveyors sampled mature forests where spatial heterogeneity was relatively low. We selected plot locations for recent surveys using original plot maps and environmental descriptions (elevation, slope, aspect), and observations in the field to maximize the match of current and historical abiotic conditions. In Mont-Mégantic, recent surveys included all plots within the current park boundary. Not all plot locations in Forillon were accessible, and plot selection for the recent surveys used the following criteria: (i) plots occurred in forest, excluding swamps or bogs; (ii) plots were accessible via <3-4 hours hiking off of established trails; (iii) plots had not obviously experienced major natural disturbances (e.g., storm, fire, or insect outbreak); (iv) in the original survey the plots were sampled in mature stands that have since maintained forest cover (i.e., no early successional dynamics in the intervening period) (Becker-Scarpitta et al. 2019).

Original surveys in Forillon Park were conducted in June-September 1972 in 256 vegetation plots of 500 m^2^ (Majcen 1981). We resurveyed 49 plots during July and August of 2015. Original surveys in Mont-Mégantic were conducted in 1970 in 94 plots, roughly half of which fall outside the current park boundaries. Plots were 400 m^2^ in coniferous forest and 800 m^2^ in broadleaved forests (Marcotte and Grandtner 1974). We resurveyed the 48 plots falling within the current park limits at Mont-Mégantic for vascular plants in 2012 (see Becker-Scarpitta et al., 2019) and for bryophytes during June and July 2014 (reported in the present study). We harmonized the taxonomy across both parks and periods (see below), so the Mont-Mégantic data are not precisely the same as reported in Savage & Vellend, (2015), but they are exactly the same as in Becker-Scarpitta et al., (2019) except transformed into presence-absence and subsetted to keep only species with at least two occurrences (see below). Raw data are stored in the *forestREplot* database (http://www.forestreplot.ugent.be) under the code: N_AM_005 for Mont-Mégantic and N_AM_008 for Forillon.

### Taxonomical database

Our taxonomical reference was the Taxonomic Name Resolution Service v4.0 (assessed in Feb 2017: http://tnrs.iplantcollaborative.org) for vascular plants and Flore des bryophytes du Québec-Labrador (Faubert, 2012, 2013, 2014) for bryophytes.

Our data set was collected by four different survey teams: one for each of the two original surveys: Forillon: Majcen, (1981); Mont-Mégantic: Marcotte & Grandtner, (1974); one for the recent Mont-Mégantic vascular plant survey: Savage & Vellend, (2015); and one for the recent Mont-Mégantic bryophyte survey and for the recent survey of both taxonomic groups at Forillon (A. Becker-Scarpitta and assistants). Most plants were identified to the species level in the same way across surveys, so for these species the only harmonization step was to standardize names. Coarser levels of taxonomic resolution were used in some but not all surveys for certain species (e.g., a pair of similar species not identified to the species level), and for other species (e.g., spring ephemeral species) the timing of different surveys created doubt about the likelihood of comparable detection. In these situations, comparability was maximized by using the coarser level of resolution applied to all data sets, or by removing species (see Supplementary material S2 for details on taxonomic standardization). All specimens identified at the species level were deposited in the Marie-Victorin herbarium (Université de Montréal, Canada.) and all locations were entered into the GBIF database (GBIF - https://www.gbif.org/).

### Statistical analysis

All statistical analyses were performed in R v.3.4.2 (R Foundation for Statistical Computing 2018). To minimize the effect of potential bias due to uncertain species identification or very rare species, all statistical analyses used the subset of species with at least two occurrences in each time period for each park.

First, to test for upward elevational shifts in species distributions along the elevational gradients, we calculated the mean elevation across occurrences for each species (Z), in each park, at each time. We modelled main effects and two- and three-way interactions between park (P), time (Y) and taxonomic group (T), each as a categorical variable. We thus tested for upward elevational shifts in species distributions, and how these shifts depend on park and taxonomic group. The model was given by:

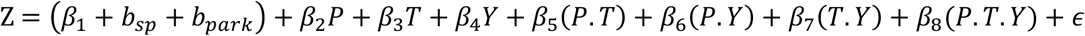

Where Z is the mean species elevation, β_1−8_ are parameters to be estimated, *b*_*sp*_ and *b*_*park*_ are random intercepts accounting for the effect of species and park, and ϵ is the model residuals assumed to be normally distributed and independent. Following our first hypothesis, we expected the temporal change at Mégantic to be stronger than at Forillon (significant *β*_*6*_ parameter). The second hypothesis predicted vascular plants to be more sensitive than bryophytes (significant *β*_*7*_ parameter). All other terms of the model that are not part of our hypotheses ensure accurate estimation of other parameters in the model.

Second, we used the exact same model structure and hypotheses to analyze temporal trends of α-diversity of both bryophytes and vascular plants in the two parks. All linear mixed effect models were conducted in R, with the function *lmer*, package ‘lme4’ v.1.1-14, Bates, Mächler, Bolker, & Walker, 2015).

Third, we tested for temporal changes in community composition using two frameworks. In one framework, we first tested the multivariate homogeneity of group dispersions - i.e. a multivariate beta-diversity metric - using the Jaccard dissimilarity index (PERMDISP, function *betadisper*, package ‘vegan’ v.2.4-4; Anderson, Ellingsen, & McArdle, (2006)). Then, we quantified the change in community composition over time using a permutational analysis of variance (PERMANOVA, Anderson, 2001) with Jaccard distances using 999 permutations to (functions *adonis*, package ‘vegan’, and *pairwise*.*adonis*, package ‘pairwiseAdonis’). We first tested the significance of the three-way interaction (park * taxonomic group * time) in a full model (F=4.45, p<0.01), and second, we ran separate models for each combination of taxonomic group and park. The R^2^ values computed from these separate PERMANOVA models were used to compare the magnitude of temporal change between groups. We used non-metric multidimensional scaling (nMDS) with Jaccard distances for visualization (function *metaMDS*, package ‘vegan’).

In a second framework, to focus on the temporal component of change in species composition, we calculated for each plot the temporal pairwise β-diversity using the binary Jaccard dissimilarity index (for presence-absence data) between original and recent surveys (function *beta*.*temp*, package ‘betapart’ v.1.5.2, Baselga & Orme, 2012). This framework allowed us to decompose the Jaccard dissimilarity into turnover i.e species replacement between plot and nestedness i.e. species loss between plot, (Baselga 2010).

As a complementary analysis, we identified for each park separately the indicator species that characterized each period using the IndVal procedure (function *multipatt*, package ‘indicspecies’; De Cáceres, Legendre, & Moretti, 2010).

## Results

### Species distributions along elevation gradients

Among the four taxonomic group-site combinations, we failed to detect any significant upward shifts in mean species elevations over time – i.e., none of the main effects or interaction terms including time were significant (Table 1a). It is important to note that patterns reported here are not quantitatively the same as in Becker-Scarpitta et al. (2019), who reported upward elevational shifts of vascular plants at Mont Mégantic based on abundance-weighted mean elevations. The difference in this paper is the use of presence-absence data, in addition to our selection of a subset of species with at least two occurrences per survey.

**Table 1 -.**
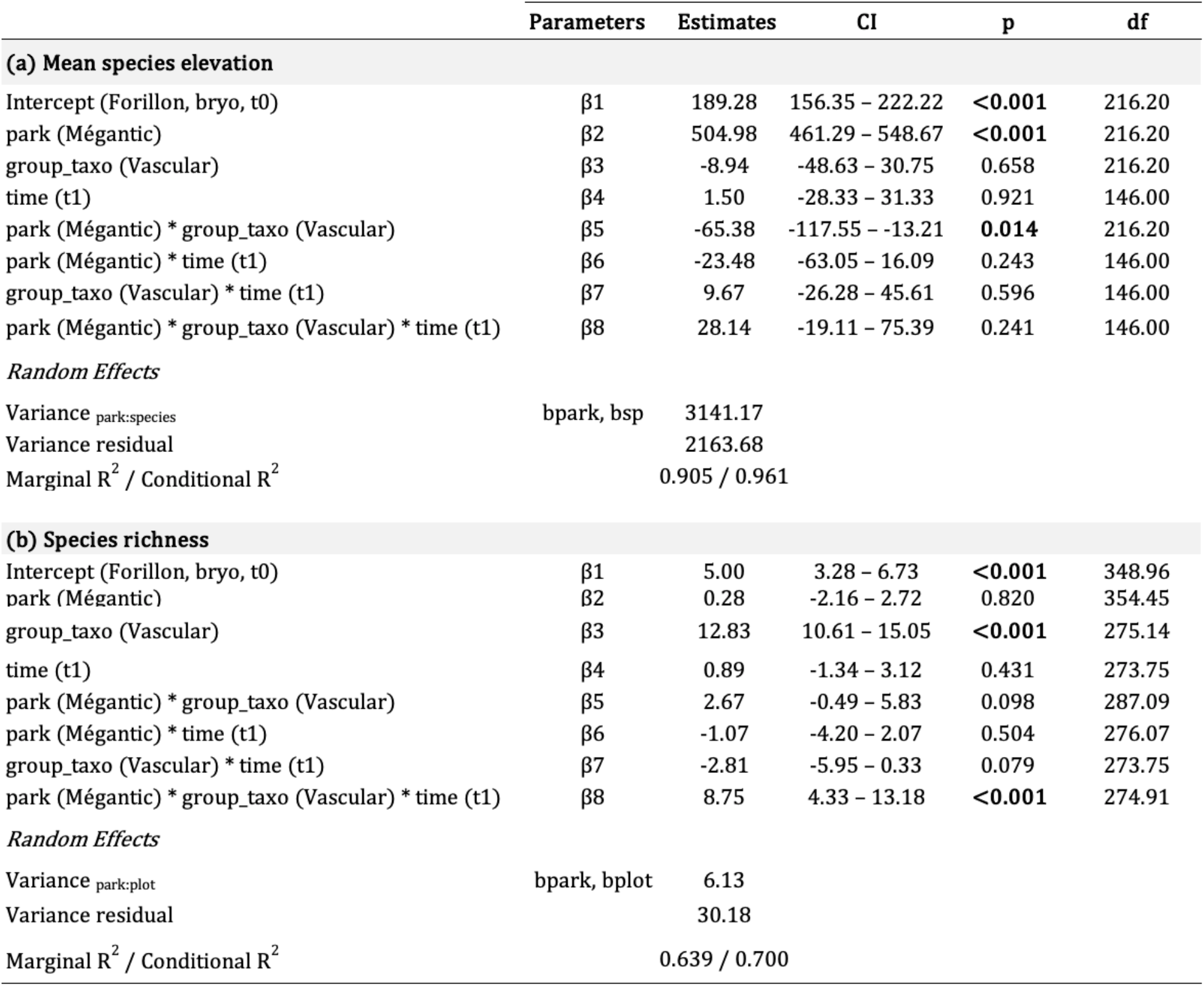
Analyses of temporal changes in (a) mean species elevation, and (b) species richness. For both models, parameter estimates ± credible intervals (CI) are reported from linear mixed models. The baseline condition (Intercept) against which effects are tested was bryophytes at Parc Forillon in the initial survey. R^2^m is the marginal coefficient of determination, measuring the proportion of variance explained by fixed effects i.e., time, park, and taxonomic group; R^2^c is the conditional coefficient of determination, giving the proportion of variance explained by both fixed and random effects. Names in parentheses refer to the level of the variable tested. In categorical models, the Intercept is the reference level used to test the effect of other variables. Bold values indicate statistically significant differences (p<0.05).

Despite the lack of trends in mean species elevation for bryophytes, there was substantial variation among species, more than for vascular plants (Table 1a, Fig. 1). This observation is reflected in the strength of correlations between original and recent mean species elevations, which was lower for bryophytes (Forillon Pearson *r* correlation = 0.33; Mont-Mégantic *r* = 0.61) than for vascular plants (Forillon *r* = 0.52; Mont-Mégantic *r* = 0.73; see Supplementary material S2 for mean elevations and sums of occurrences for each species in each survey).

**Figure 1.**
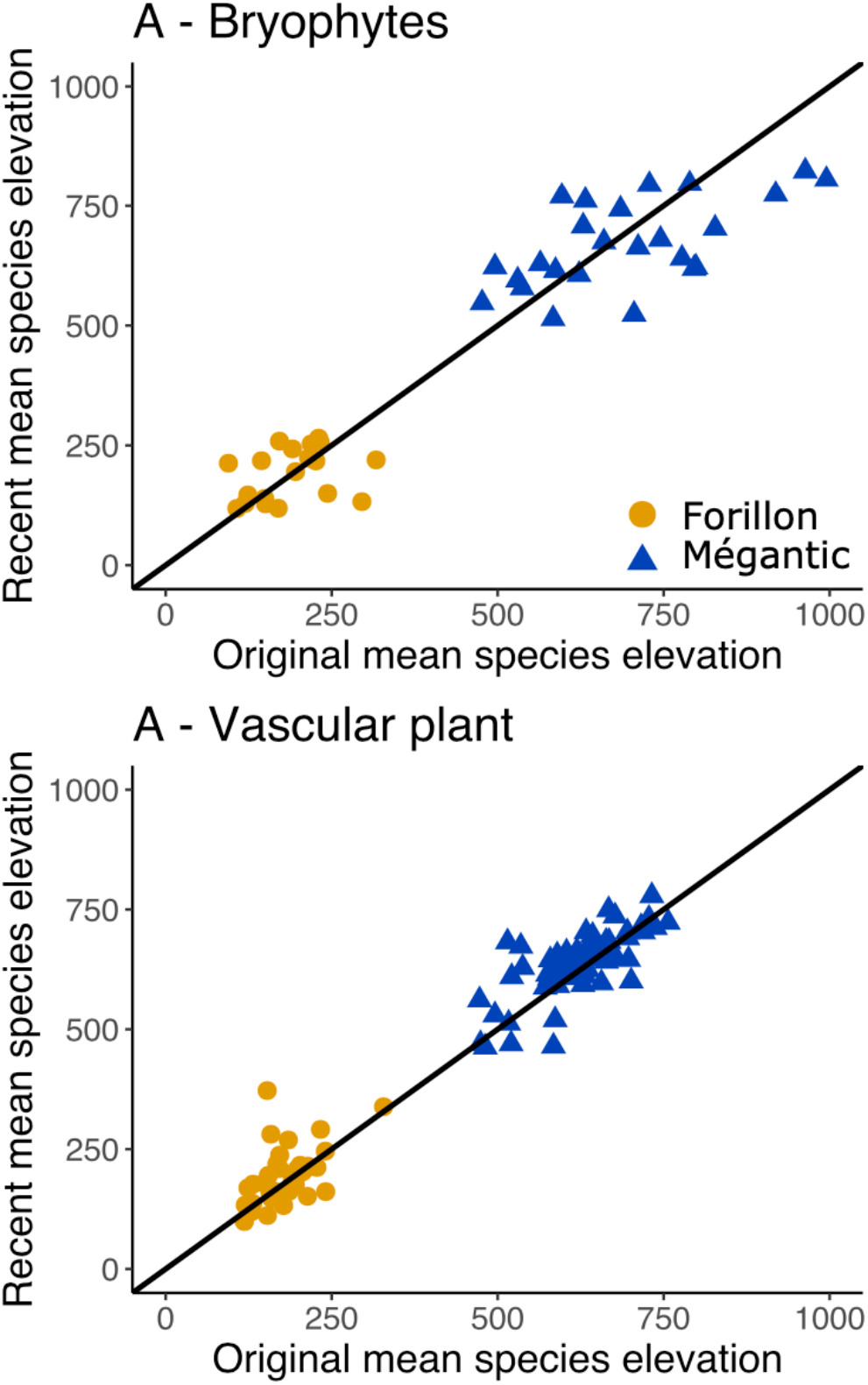
Mean elevations of species’ distributions in two time periods and two parks. A) Bryophytes, Forillon = 19 species, Mégantic = 25 species); B) Vascular plants, Forillon = 42 species, Mégantic = 64 species. All species present in both the original and recent surveys were included. Diagonal lines indicate 1:1 (i.e. no changes over time). See Table 1a for statistical analysis.

### Temporal changes in species richness

At Forillon, the total number of bryophyte species across plots was greater in the recent survey (38 species) than in the original survey (25 species); 6 species from the original survey were not found while 19 new species were observed, and the two surveys shared 19 species (Supplementary material S3 for species occurrences). Similarly, at Mont-Mégantic, there was an increase in the overall number of bryophytes species in the recent survey (original survey = 31 vs. recent = 39 species), with 6 species lost, 14 gained while 25 species were shared between the two surveys.

Conversely, the total number of vascular plants at Forillon declined over time (original survey = 64 vs. recent = 44 species), with 22 original species not found in the recent survey, 2 new species added and 42 shared species. While at Mont-Mégantic, the total number of vascular plant species increased (from 72 species to 77 species), with 8 species lost, 13 gained and 64 shared species.

We detected a significant increase in mean plot-level species richness (α-diversity) only for vascular plants at Mont-Mégantic (Table 1b). For bryophytes in both parks, and for vascular plants at Forillon α-diversity showed no changes over time (Table 1b).

### Temporal shift in community composition and heterogeneity

We found significant temporal changes in vascular plant β-diversity and community composition at Mont-Mégantic (Table 2, Fig. 2), where climate warming has been pronounced, as previously reported (Becker-Scarpitta et al. 2019). However, we found an unexpected significant increase of multivariate dispersion (β-diversity) over the past 40 years for bryophytes at Forillon but not at Mont-Mégantic (Table 2, Fig. 2).

**Table 2.** Temporal changes in beta diversity and community composition. β-diversity was measured as the multivariate distance of plots to the time-specific centroid in multivariate space. The R^2^ of the shift in community composition reflects the proportion of variation in community composition explained by time. Jaccard temporal β-diversity values are decomposed into turnover and nestedness between original and recent surveys. Bold values indicate significant statistical differences p<0.05.

| | | β-diversity | | | Composition | | | Temporal beta diversity | | |
| --- | --- | --- | --- | --- | --- | --- | --- | --- | --- | --- |
|  |  | Original | Recent | p | F | p.adj | R2 | Jaccard | Turnover(%) | Nestedness(%) |
| Bryophytes |  |  |  |  |  |  |  |  |  |  |
|  | Forillon | 0.53 | 0.58 | **0.015** | 6.93 | **0.006** | 0.070 | 0.80±0.02 | 85.5% | 14.5% |
|  | Megantic | 0.61 | 0.60 | 0.93 | 7.64 | **0.006** | 0.074 | 0.89±0.02 | 96.4% | 3.6% |
| Vascular plants |  |  |  |  |  |  |  |  |  |  |
|  | Forillon | 0.45 | 0.43 | 0.80 | 1.86 | 0.126 | 0.019 | 0.52±0.02 | 69.9% | 29.9% |
|  | Megantic | 0.50 | 0.43 | **0.004** | 3.60 | **0.024** | 0.037 | 0.53±0.02 | 70.8% | 29.2% |

**Figure 2.**
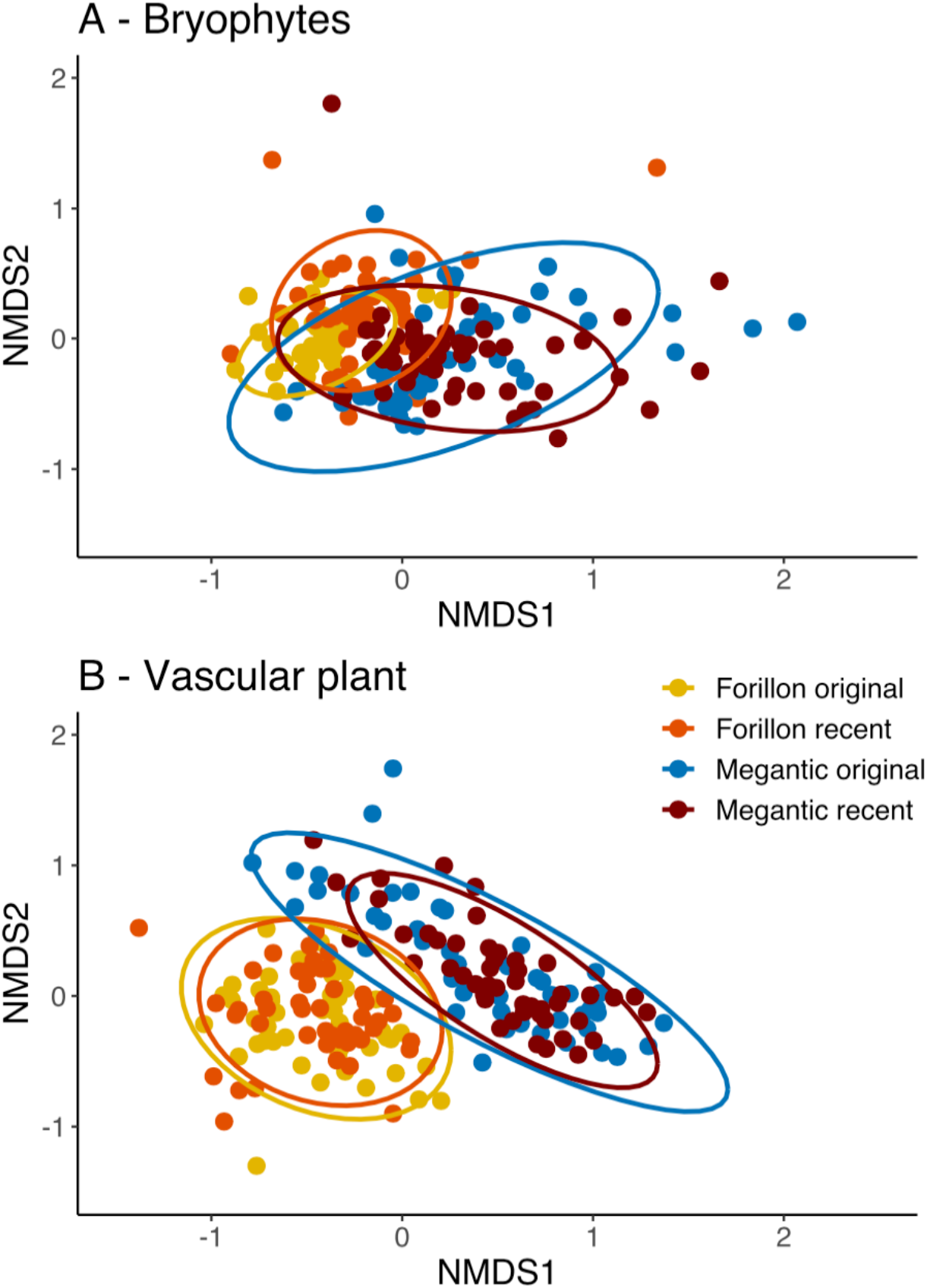
Temporal changes in (A) bryophyte and (B) vascular plant community composition over time in Forillon and Mont-Mégantic, based on Jaccard’s dissimilarity and non-metric multidimensional scaling (NMDS). At both sites, bryophytes showed significant shifts in composition over time and a significant increase in beta diversity. For vascular plants significant shifts in community composition and a significant decrease in beta diversity were found at Mont Mégantic but not Forillon (Table 2).

For both sites we found significant temporal shifts in bryophyte community composition; for vascular plants there was significant compositional change only at Mont-Mégantic (Table 2, Fig. 2). As predicted, the magnitude of the vascular plant compositional shift was greater for Mont-Mégantic (R^2^=0.037) than Forillon (R^2^=0.019). However, bryophyte communities experienced an equal magnitude of compositional shift for both Forillon and Mont-Mégantic (R^2^=0.07 and 0.074 respectively). Moreover, compositional shifts were greater for bryophytes than for vascular plants at both sites (Table 2).

Temporal beta-diversity showed contrasted patterns between taxonomic groups. Jaccard dissimilarity between original and recent surveys was higher for bryophytes than for vascular plants. The greater change in temporal beta diversity for bryophytes in the two parks is consistent with the PERMANOVA results for changes in community composition. For all combinations of taxonomic group and park, species turnover contributed more than nestedness to overall changes. However, nestedness was considerably higher for vascular plants than for bryophytes (Table 2). In support of the beta-diversity analysis, Table 3 shows the indicator species list that characterized each time period in each park. All combinations of park and time show a complete change in the indicator species assemblage. Even in the absence of beta-diversity change for bryophytes at Mégantic or for vascular plants at Forillon, we detected changes in certain species frequencies. Several species associated with recent time periods suggest possible non-climatic drivers of community change (e.g., Dennstaedtia punctilobula and Carex spp. are resistant to deer herbivory), a point we develop further in the discussion.

**Table 3 -.**
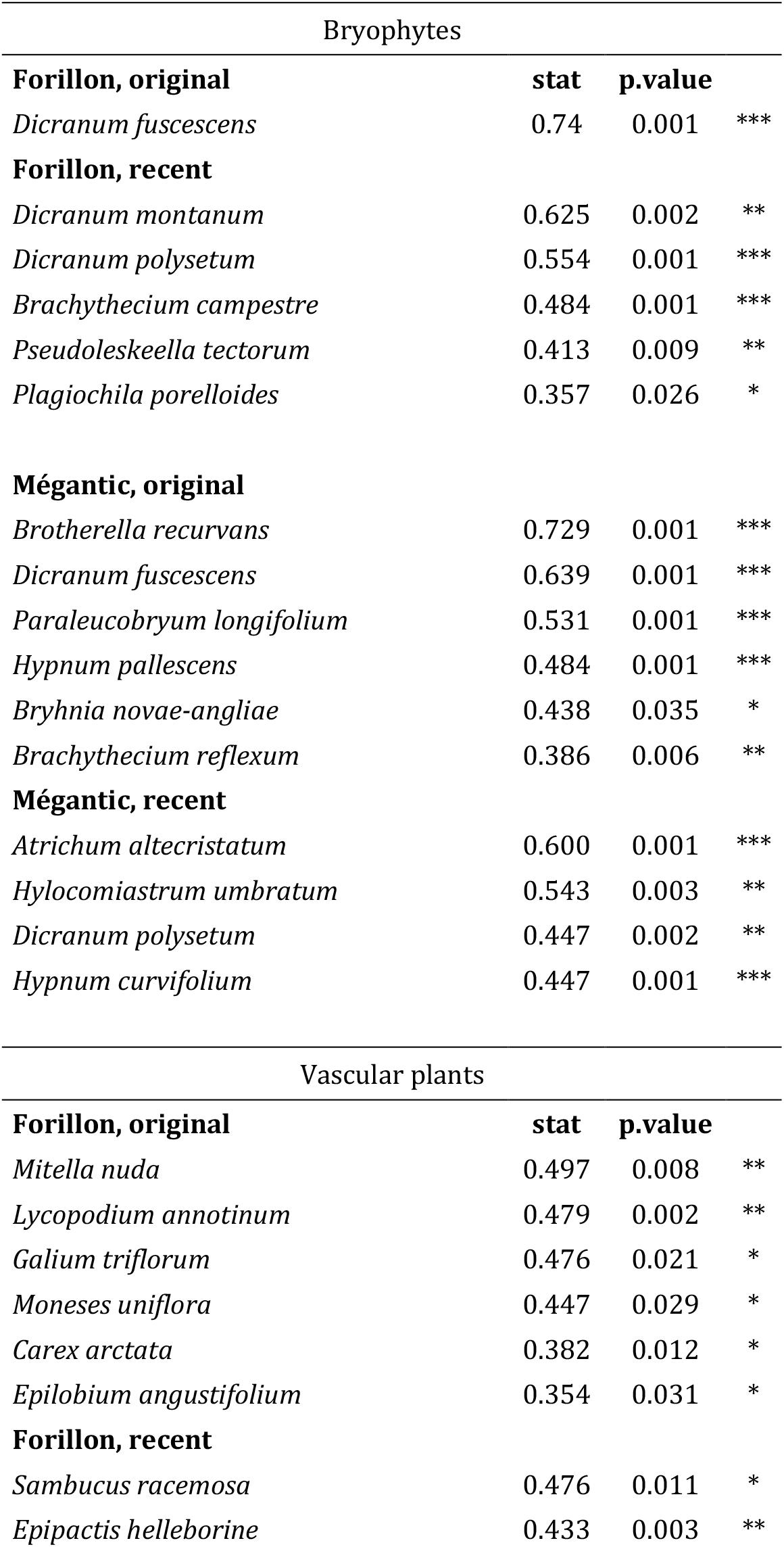
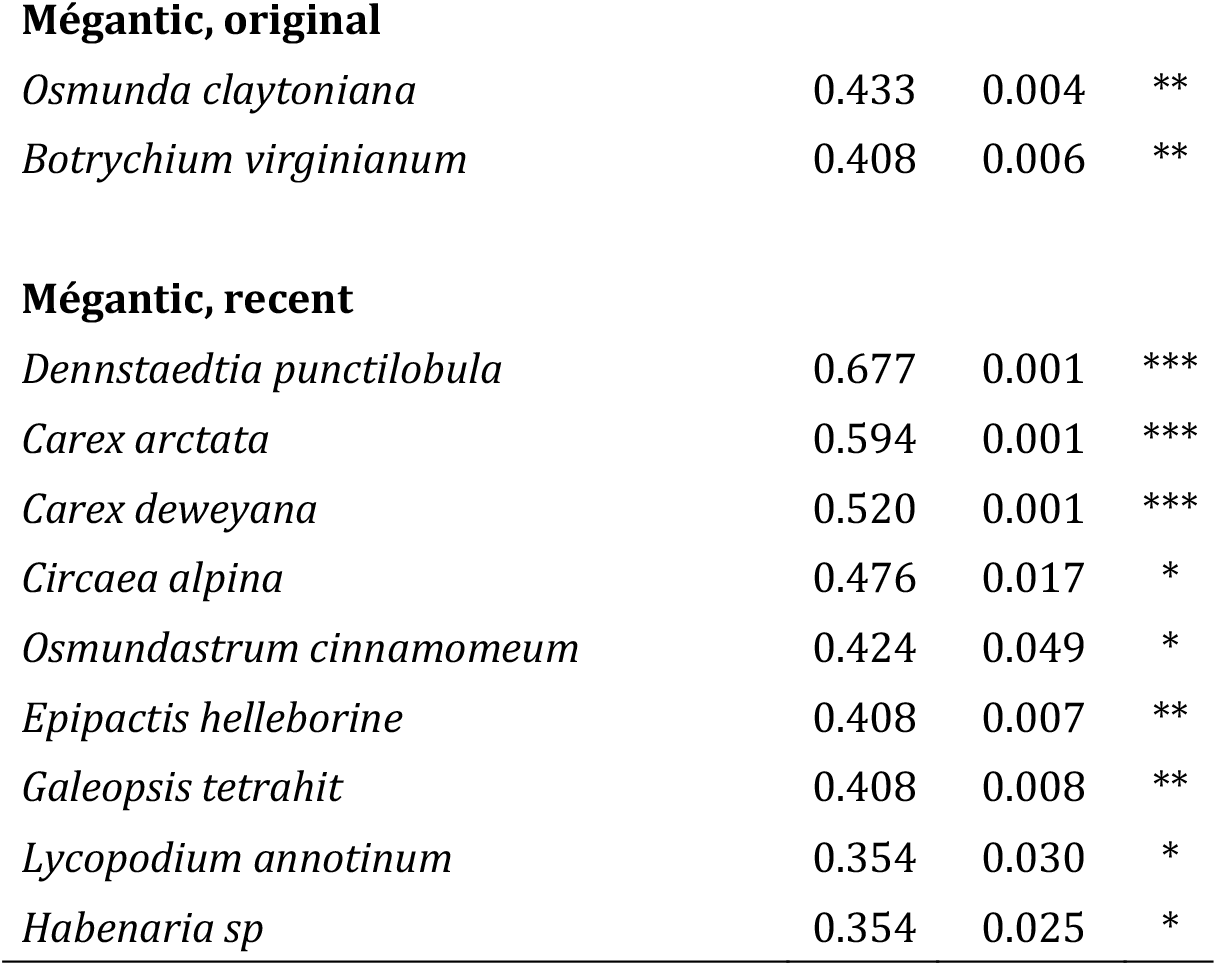
Indicator species of bryophytes and vascular plants associated with original and recent surveys at Forillon and Mont-Mégantic. Indicator values were considered significant if p<0.05 (calculated with 999 permutations).

| Bryophytes |  |  |  |
| --- | --- | --- | --- |
| **Forillon, original** | **stat** | **p.value** |  |
| *Dicranum fuscescens* | 0.74 | 0.001 | *** |
| **Forillon, recent** |  |  |  |
| *Dicranum montanum* | 0.625 | 0.002 | ** |
| *Dicranum polysetum* | 0.554 | 0.001 | *** |
| *Brachythecium campestre* | 0.484 | 0.001 | *** |
| *Pseudoleskeella tectorum* | 0.413 | 0.009 | ** |
| *Plagiochila porelloides* | 0.357 | 0.026 | * |
| **Mégantic, original** |  |  |  |
| *Brotherella recurvans* | 0.729 | 0.001 | *** |
| *Dicranum fuscescens* | 0.639 | 0.001 | *** |
| *Paraleucobryum longifolium* | 0.531 | 0.001 | *** |
| *Hypnum pallescens* | 0.484 | 0.001 | *** |
| *Bryhnia novae-angliae* | 0.438 | 0.035 | * |
| *Brachythecium reflexum* | 0.386 | 0.006 | ** |
| **Mégantic, recent** |  |  |  |
| *Atrichum altecristatum* | 0.600 | 0.001 | *** |
| *Hylocomiastrum umbratum* | 0.543 | 0.003 | ** |
| *Dicranum polysetum* | 0.447 | 0.002 | ** |
| *Hypnum curvifolium* | 0.447 | 0.001 | *** |
| Vascular plants |  |  |  |
| **Forillon, original** | **stat** | **p.value** |  |
| *Mitella nuda* | 0.497 | 0.008 | ** |
| *Lycopodium annotinum* | 0.479 | 0.002 | ** |
| *Galium triflorum* | 0.476 | 0.021 | * |
| *Moneses uniflora* | 0.447 | 0.029 | * |
| *Carex arctata* | 0.382 | 0.012 | * |
| *Epilobium angustifolium* | 0.354 | 0.031 | * |
| **Forillon, recent** |  |  |  |
| *Sambucus racemosa* | 0.476 | 0.011 | * |
| *Epipactis helleborine* | 0.433 | 0.003 | ** |
| **Mégantic, original** |  |  |  |
| *Osmunda claytoniana* | 0.433 | 0.004 | ** |
| *Botrychium virginianum* | 0.408 | 0.006 | ** |
| **Mégantic, recent** |  |  |  |
| *Dennstaedtia punctilobula* | 0.677 | 0.001 | *** |
| *Carex arctata* | 0.594 | 0.001 | *** |
| *Carex deweyana* | 0.520 | 0.001 | *** |
| *Circaea alpina* | 0.476 | 0.017 | * |
| *Osmundastrum cinnamomeum* | 0.424 | 0.049 | * |
| *Epipactis helleborine* | 0.408 | 0.007 | ** |
| *Galeopsis tetrahit* | 0.408 | 0.008 | ** |
| *Lycopodium annotinum* | 0.354 | 0.030 | * |
| *Habenaria sp* | 0.354 | 0.025 | * |

## Discussion

Most long-term legacy studies have been conducted on vascular plant communities (Verheyen et al. 2017), so it remains unclear whether bryophytes show similar or different responses to environmental changes. Here we quantified community changes over ∼40 years for both bryophytes and vascular plants in two protected natural areas in eastern Canada with contrasting recent warming trends. Temporal changes in vascular plant communities contrasted with the warming hypothesis. As predicted for the area with a strong warming trend (Mont-Mégantic), we found a significant increase of local diversity of vascular plants but no change for bryophytes (Table 1b), and a stronger change in β-diversity and community composition for vascular plans as well (Table 2). However, we did not detect any shift in species distribution along the elevation gradient (Table 1a, Fig. 1) and the higher magnitude of changes in bryophyte community composition at both sites was not expected (Table 2, Fig. 2).

### Patterns along the gradient of warming trends

At Mont-Mégantic, where the warming trend has been strongest, we found a significant increase of α-diversity (Table 1b), a significant decrease of β-diversity and a stronger shift in community composition than in Forillon (Table 2, Fig. 2). At Forillon, where the warming trend has been weaker, we found neither shifts in elevational distributions (Table 1a, Fig. 1) nor changes of α-diversity for either vascular plants or bryophytes (Table 1a). The lack of upward shift in elevation of vascular plants in response to warming is clearly explained by the use of presence/absence data, and contrasts with our previous results using abundance data (Becker-Scarpitta et al. 2019) in addition to studies from other sites (Lenoir et al. 2008, Chen et al. 2011, Bertrand et al. 2011),. The temporal increase of diversity of vascular plants at Mont-Mégantic is also consistent with the prediction that warming should lead to increased local diversity in areas without severe moisture stress (Vellend et al. 2017b, Harrison 2020).

Our results suggest that a temperature increase of ∼0.9 °C (over the last 45 years at Mont-Mégantic) does not have as strong an impact on the local diversity of bryophytes as it does for vascular plants, assuming that the richness increase at Mégantic was indeed due to warming. This interpretation is also supported by the absence of any relationship between bryophyte richness and elevation in the two parks (Forillon estimate= 0.0042 ± (SE) 0.0024, t=1.8, p=0.079 ; Mégantic estimate= 0.0022 ± (SE) 0.0014, t=1.6, p= 0.11) and consistent with previous results in the literature (Bruun et al., 2006; Grytnes, Heegaard, & Ihlen, 2006; Odland, Reinhardt, & Pedersen, 2014 but see Vittoz et al., 2010). The decrease of β-diversity over time for vascular plants at Mont-Mégantic is consistent with the hypothesis that warming might cause biotic homogenization (Urban 2015, Socolar et al. 2016). Note that this result using presence-absence data was different to the result in Becker-Scarpitta et al., (2019) using untransformed abundance data (for which we found no homogenization), but similar to the finding of homogenization using fourth-root transformed abundances in Savage & Vellend, (2015). These results collectively illustrate that the locally dominant species – whose influence is minimized or eliminated via fourth-root or presence-absence transformation – can mask homogenization created by species of lower abundance. This result highlights how ecological conclusions are dependent on the data transformation, even with widely used diversity metrics.

### Sensitivity of bryophytes vs. vascular plants

We cannot draw strong conclusions about which of bryophytes or vascular plants is more or less sensitive to environmental change in eastern Canada: results depended on which community dimension was being investigated. Vascular plants showed more prominent richness increases (Tables 1b) and a decrease in β-diversity, while bryophytes experienced stronger shifts in community composition and no difference in the magnitude of compositional changes between the two parks (Table 2, Fig. 2). We did not detect (with presence-absence data) shifts in distribution along the elevational gradient for either taxonomic group (Table 1a, Fig. 1), while we previously found a clear shift in elevation when using vascular plant abundance data at Mont-Mégantic (Becker-Scarpitta et al. 2019). Our results for elevational shifts here are thus inconclusive, in that we do not know if analyses using abundance data for bryophytes would have also shown a change in elevation.

### Potential non-climatic drivers of vegetation change

Our sites were chosen specifically due to their contrasting warming trends and lack of other obvious major drivers of vegetation change, but there are certainly other possible drivers of ecological change that might play a role in this region. Among the indicator species of recent surveys (Table 3), two were non-native: *Galeopsis tetrahit* and *Epipactis helleborine*, the latter of which has increased considerably in recent decades throughout its North American range, even in western Canada (Marie-Victorin 1997, McCune and Vellend 2013). As such, some vegetation changes might be due simply to protracted periods of non-native species expansions, regardless of local environmental change. Another potential factor is changes in white-tailed deer browsing, which has increased over the past century in much of North America (Côté et al. 2004). The indicator species of the recent survey at Mont-Mégantic (Table 3) include species known to benefit from high levels of deer browsing: *Dennstaedtia punctilobula* and *Carex* spp. (de la Cretaz and Kelty 2002, Augustine and Decalesta 2003, Rooney 2009, Frerker et al. 2014). Interestingly, at Forillon deer are actually thought to have decreased in abundance due to the expansion of the coyote population in the 1970s (UQCN 2005), and we found no such species associated with recent surveys in Forillon Park.

Our most difficult result to interpret was the strong species turnover of bryophyte communities at Forillon, which has not experienced strong long-term increase in temperature, precipitation, or atmospheric nutrient deposition (Commission Joint International 2014, Hember 2018). We can only speculate regarding potential non-climatic drivers of bryophyte community change. First, as in all legacy studies, there is the potential for observer biases due to (i) different sampling effort between original and recent surveys, or (ii) species’ identification errors. It seems highly likely that detection probabilities and the potential for identification errors are greater for bryophytes than for vascular plants, although we have no reason to suspect this caused systematic increases or decreases of particular species frequencies (necessary to explain overall compositional shifts). We paid very close attention to repeating the original survey methods, focusing on the visually obvious species in each microsite (i.e., not examining each individual moss stem closely on the field), and the lack of any difference over time in local richness suggests comparable species’ detection abilities in the two surveys. Although we cannot exclude the possibility that a real richness change was cancelled out by a change in survey effort, we have no reason to suspect this rather unlikely coincidence. Second, given uncertainty in the comparability of abundance estimates across time for bryophytes, we also decided to use presence-absence data. In short, changes in observer effort seem highly unlikely to account for the temporal change in species composition. Furthermore, our procedure to homogenize taxonomy across data sets was quite conservative to reduce bias due to misidentifications.

One potential hypothesis to explain compositional change over time in bryophyte communities relates to the history of park protection. Forillon Park was established (and so protected) only two years before the original survey was conducted, and parts of the park previously included homesteads (i.e., non-forest land uses). Mont-Mégantic was established as a park more recently (1994), but logging activities (the only prominent land use) ceased ∼15 years before the original survey. Although plot selection focused only on non-disturbed forests, metacommunity dynamics involving dispersal of species from sites undergoing rapid succession may have caused local shifts in composition and increased β-diversity. It has been previously shown that managed forests tend to have a lower β-diversity than protected forests (Kaufmann et al. 2017). The increase in bryophyte β-diversity might partially be due to the recovery of natural forest that occurring in the 1970s.

There is also the possibility that changes in bryophyte communities resulted from interactions with changing vascular plant communities. Studies have shown that bryophyte diversity and abundance can be negatively correlated with total vascular plant biomass (Virtanen et al. 2017), cover (Jiang et al. 2015) or abundances (Jägerbrand et al. 2012). While we have documented an increase of vascular plant species richness at Mont-Mégantic (Tables 1b), we do not have data on vascular plant biomass. If bryophytes are highly sensitive to vascular plant community properties, subtle changes for vascular plants could translate into larger changes in bryophyte communities. This hypothesis is open to testing via observational and experimental studies of the effect of vascular plants on bryophytes communities under warming or other environmental changes. Understanding temporal changes in one component of the community may require more consideration of interactions with other components (Chesson 2000, HilleRisLambers et al. 2012).

### Conclusion and conservation implications

Overall, we found a significant temporal shift in the composition of both taxonomic groups in both parks but only one significant change in α-diversity. This study shows the limitation of using presence absence data to detect shift in distribution. While abundance data provide the necessary power to detect changes in species distributions along the elevational gradient (Becker-Scarpitta et al. 2019), presence/absence data clearly do not. Our results are in accordance with recent meta-analyses and syntheses showing that local diversity can remain unchanged (or increase or decrease with equal likelihood) despite strong changes in composition (Dornelas et al. 2014, Vellend et al. 2017a, Gotelli et al. 2017, Spaak et al. 2017, Magurran et al. 2018, Blowes et al. 2019). Finally, regardless of whether one taxonomic group is systematically more or less sensitive to environmental change than another, our results suggest that one taxonomic group (e.g., vascular plants) cannot be used as a surrogate for others (e.g., bryophytes) in terms of predicting the nature and magnitude of responses to environmental change (Bagella 2014, Becker-Scarpitta et al. 2017, Outhwaite et al. 2020). In the same plots that experienced the same environmental changes, we found that communities of bryophytes and vascular plants did not predictably change in the same ways (Slack 1977, Lalanne et al. 2008, Odland et al. 2014, Bagella 2014, Becker-Scarpitta et al. 2017). Thus, to assess overall biodiversity responses to global change abundance data from different taxonomical groups and different community properties need to be synthesized.

## Supporting information

Supplementary_S1_climate

Supplementary_S2_taxonomy

Supplementary_S3_elev_occurrence

Supplementary_S4_IndVal

## Acknowledgements

We would like to thank the field assistants who greatly contributed to the collection of data in the field, the identification of species in the laboratory and the realization of the herbarium: Melissa Paquette and Sara Gaignard. This work was made possible thanks to the support of park managers, especially Camille-Antoine Ouimet (at Mont-Mégantic) and Daniel Sigouin (at Forillon). We also thank Guillaume Blanchet, Michael Belluau and Willian Vieira for constructive discussion on this project. The project was funds by the Natural Sciences and Engineering Research Council, Canada and field mission were support by the Quebec Center for Biodiversity Sciences (QCBS).

## Data Accessibilty Statement

Data are stored in the *forestREplot* database (http://forestreplot.ugent.be/) under the code: N_AM_005 for Mont-Mégantic and N_AM_008 for Forillon. All plant specimens identified at the species level were deposited in the Marie-Victorin herbarium (Université de Montréal, Canada.) and all locations were entered into the GBIF database (GBIF - https://www.gbif.org/).

## Supplementary Materials

- Appendix S1 – Climatic temporal trends of the two parks.
- Supplementary material S2 – Taxonomic standardization of bryophytes species
- Supplementary material S3 – Species occurrences and mean species elevations

## Appendix S1

### Climatic temporal trends

To model the mean annual temperature trend in both parks, we extracted temperature data from the ANUSPLINE model (McKenney et al. 2011). The relevant period to study the effect of temperature change on forest plant communities includes a lag of approximately ∼10 years before the time of the survey. We thus considered the period 1960-2005, which aligns with a previous study showing a sharp warming gradient in the region (Yagouti et al. 2008). The model includes the effect of the categorical variable park (with two levels: Forillon and Mégantic), the continuous variable year from 1960 to 2005, and a random effect on the different measures within each park.

We only report results for annual mean temperature but found similar results for annual minimum and maximum temperatures, with no temporal change in annual mean precipitation. For the period 1960-2005, we find that Forillon experienced an increase of 0.12 °C ± 0.010°C/decade, while at Mégantic the increase per decade was almost twice as strong: 0.20 °C ± 0.014°C/decade.

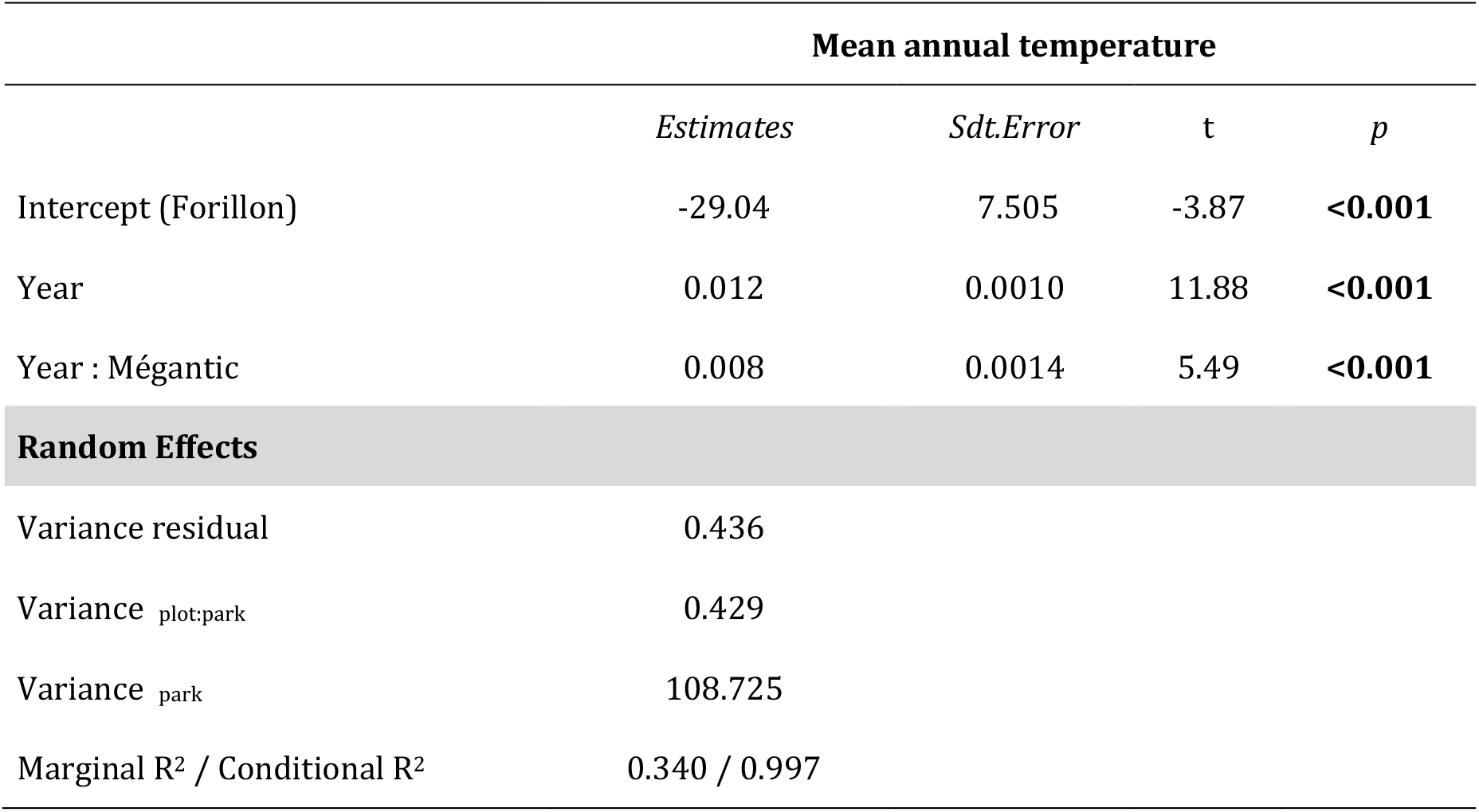

