## Supplementary_S1_climate for "Different temporal trends in vascular plant and bryophyte communities along elevational gradients over four decades of warming"

**Appendix S1 : Climatic temporal trends**

To model the mean annual temperature trend in both parks, we extracted temperature data from the ANUSPLINE model (McKenney et al. 2011). The relevant period to study the effect of temperature change on forest plant communities includes a lag of approximately ~10 years before the time of the survey. We then consider the period 1960-2005, which aligns with a previous study showing a sharp warming gradient in the region (Yagouti et al. 2008). The model includes the effect of the categorical variable park (with two levels: Forillon and Megantic), the continuous variable year from 1960 to 2005 and a random effect on the different measures within each park.

We only reported results for annual mean temperature, but found similar results for annual minimum and maximum temperatures, with no temporal change in annual mean precipitation.For the more relevant period 1960-2005, we find that Forillon experiences an increase of 0.12 ^o^C ± 0.010^o^C/decade, while at Megantic the increase per decade is twice as great: 0.20 ^o^C ± 0.014^o^C/decade.

|  | **Mean annual temperature** | | | |
| --- | --- | --- | --- | --- |
|  | *Estimates* | *Sdt.Error* | t | *p* |
| Intercept (Forillon) | -29.04 | 7.505 | -3.87 | **<0.001** |
| Year | 0.012 | 0.0010 | 11.88 | **<0.001** |
| Year : Megantic | 0.008 | 0.0014 | 5.49 | **<0.001** |
| **Random Effects** |  |  |  |  |
| Variance residual | 0.436 |  |  |  |
| Variance  _plot:park_ | 0.429 |  |  |  |
| Variance  _park_ | 108.725 |  |  |  |
| Marginal R^2^ / Conditional R^2^ | 0.340 / 0.997 |  |  |  |
