## Supplementary_S2_taxonomy for "Different temporal trends in vascular plant and bryophyte communities along elevational gradients over four decades of warming"

**Appendix S2 – Taxonomic standardization of bryophytes species**

We present here a detailed account of taxonomic standardization of bryophytes species across different data sets (four surveys made by 3 different field teams). As a rule, any decision made by a given survey (e.g., merging species because of high morphological resemblances) was applied to all surveys. Homogenization of data sets involved the following merging procedure:

- In cases for which one species in the 1970s has since been split into multiple species, we use the original species name even if it now refers to a complex of species.
- Species with very close morphologically sharing the same ecological requirements were merge under a single name.

|  | **Doubtfull species** | **Merge into** |
| --- | --- | --- |
| **Forillon original** |  |  |
|  | *Hypnum lindbergii* | *Hypnum lindbergii* |
|  | *Hypnum pratense* |  |
|  | *Hylocomium pyrenaicum* | *Hylocomiastrum umbratum* |
|  | *Hylocomium umbratum* |  |
| **Forillon recent** |  |  |
|  | *Plagiothecium cavifolium* | *Plagiothecium cavifolium* |
|  | *Plagiothecium latebricola* |  |
|  | *Brachythecium curtum* | *Brachythecium curtum* |
|  | *Brachythecium rivulare* |  |
|  | *Brachythecium starkei* |  |
|  | *Kinbergia praelonga* |  |
|  | *Brachythecium campestre* | *Brachythecium campestre* |
|  | *Brachythecium falcatum* |  |
|  | *Brachythecium rutabulum* |  |
|  | *Barbilophozia barbata* | *Barbilophozia hatcheri* |
|  | *Barbilophozia hatcheri* |  |
|  | *Barbilophozia lycopodioides* |  |
|  | *Dicranum brevifolium* | *Dicranum fuscescens* |
|  | *Dicranum fuscescens* |  |
|  | *Herzogiella striatella* | *Herzogiella striatella* |
|  | *Herzogiella turfacea* |  |
|  | *Thuidium delicatum* | *Thuidium delicatum* |
|  | *Thuidium recognitum* |  |
| **Mégantic original** |  |  |
|  | *Amblystegium varium* | *Hygroamblystegium varium* |
|  | *Atrichum oerstedianum* | *Atrichum crispulum* |
|  | *Brachythecium rutabulum* | *Brachythecium campestre* |
|  | *Brachythecium salebrosum* | *Brachythecium acutum* |
|  | *Brachythecium starkei* | *Brachythecium curtum* |
|  | *Eurhynchium pulchellum* | *Eurhynchiastrum pulchellum* |
|  | *Hylocomium umbratum* | *Hylocomiastrum umbratum* |
|  | *Hypnum pratense* | *Hypnum lindbergii* |
|  | *Hypnum reptile* | *Hypnum pallescens* |
|  | *Isopterygium distichaceum* | *Pseudotaxiphyllum distichaceum* |
|  | *Jungermannia lanceolata* | *Jungermania leiantha* |
|  | *Lophozia attenuata* | *Barbilophozia attenuata* |
|  | *Mnium ciliare* | *Plagiomnium ciliare* |
|  | *Mnium cusidatum* | *Plagiomnium cuspidatum* |
|  | *Mnium medium* | *Plagiomnium medium* |
|  | *Mnium ponctuatum* | *Rhizomnium punctatum* |
|  | *Polytrichum gracile* | *Polytrichum longisetum* |
|  | *Polytrichum ohioense* | *Polytrichastrum pallidisetum* |
|  | *Porella platyphylloïdea* | *Porella platyphylla* |
|  | *Sphagnum centrale* | *Sphagnum sp* |
|  | *Sphagnum girgensohnii* |  |
|  | *Sphagnum recurvum* |  |
|  | *Sphagnum robustum* |  |
|  | *Sphagnum squarrosum* |  |
|  | *Sphagnum warnstorfianum* |  |
| **Mégantic recent** |  |  |
|  | *Brachythecium curtum* | *Brachythecium curtum* |
|  | *Brachythecium rivulare* |  |
|  | *Kinbergia praelonga* |  |
|  | *Brachythecium rutabulum* | *Brachythecium campestre* |
|  | *Calypogeia neesiana* | *Calypogeia neesiana* |
|  | *Calypogeia mulleriana* |  |
|  | *Dicranum ontariense* | *Dicranum polysetum* |
|  | *Dicranum polysetum* |  |
|  | *Hylocomiastrum pyrenaicum* | *Hylocomiastrum umbratum* |
|  | *Hylocomiastrum umbratum* |  |
|  | *Polytrichastrum ohioense* | *Polytrichastrum pallidisetum* |
|  | *Polytrichastrum pallidisetum* |  |
|  | *Fissidens dubius* | *Fissidens osmundoides* |
